# Generalist malaria parasites and host imprinting: Unveiling transcriptional memory

**DOI:** 10.1101/2025.03.24.644894

**Authors:** Luz García-Longoria, Vaidas Palinauskas, Juste Aželytė, Alfonso Marzal, David Ovelleiro, Olof Hellgren

## Abstract

Generalist parasites must rapidly adapt to diverse host environments to ensure their survival and transmission. Parasites may employ fixed genetic responses, transcriptional plasticity, or epigenetic mechanisms to optimize survival. The avian malaria parasite *Plasmodium homocircumflexum* serves as an ideal model for studying transcriptional variation and adaptive strategies. We experimentally inoculated *P. homocircumflexum* into different bird hosts, bypassing vector recombination, to investigate whether parasite gene expression remains stable across hosts, resets in response to new environments, or reflects epigenetic inheritance. Our study evaluates four potential mechanisms: (1) A universal gene expression profile (“one key fits all”), where expression remains stable across hosts. Our outcomes revealed that gene expression differed significantly depending on the host species and time post-infection, thus rejecting this hypothesis. (2) Complete transcriptional plasticity, where gene expression is fully determined by the recipient host. Contrary to this hypothesis, we observed that gene expression was primarily influenced by the donor at 8 days post-infection (dpi), whereas gene expression was more aligned with the recipient host at 16 dpi. (3) Epigenetic inheritance, where early-stage gene expression reflects the donor host but gradually adjusts to the recipient. Our results support this mechanism, as 2,647 differentially expressed genes (DEGs) were associated with donors at 8 dpi, whereas 271 DEGs were linked to the recipient by 16 dpi. (4) Selection-driven differentiation favoring specific haplotypes. This latter hypothesis was not supported since SNP analyses showed low genetic differentiation. These findings suggest a *P. homocircumflexum* transition from donor-dependent to recipient-dependent gene expression, likely mediated by epigenetic regulation and transcriptional plasticity.

## Introduction

The survival of generalist parasites depends on their ability to adapt to new environments (1). When conducting host shifts the parasite get exposed to varying immune responses, body temperatures, metabolism, and nutrient availability, all affecting survival and replication (2,3). Vector transmitted parasites such as *Plasmodium* spp. must navigate in both vertebrate and invertebrate hosts, with the vertebrate immune system being a major challenge (4,5). Just considering the vertebrate immune responses, they might further vary across species (6,7) and geographic regions (8,9), requiring parasites to be highly flexible to overcome these challenges. One way to achieve this flexibility is by relying on transcriptional variation, where gene expression shifts in response to environmental changes, often regulated by epigenetic mechanisms like DNA methylation (10) and histone modifications (11). These mechanisms enable adaptation without altering the genome (12), helping parasites to evade host immunity and adjust to host metabolism within a single infection.

A key question is whether generalist parasites adapt through fixed genetic responses or flexible adjustments like epigenetic modifications and real-time gene expression shifts. One possibility is a “universal key” strategy, where parasites maintain a static gene expression profile that enables infection across different hosts with minimal adjustment, relying on conserved gene responses (13,14). Alternatively, selection-driven variation may favor specific haplotypes with expression profiles better suited to certain hosts (15). Another strategy involves dynamic responses, in which gene expression adapts to different host immune defenses and metabolism (16). This plasticity may arise from epigenetic mechanisms or gene expression patterns inherited asexually from the parent population (17), enabling the replication of successful adaptive responses. These inherited profiles may shift when environmental changes demand adaptation, optimizing parasite survival across hosts (18).

*Plasmodium falciparum* exemplifies transcriptional variation, with gene expression changes regulated through complex epigenetic mechanisms (19). Genes involved in antigenic variation and immune evasion exhibit clonal variation, meaning genetically identical parasites express different gene sets (20). This allows *P. falciparum* to maintain a diverse population, some better suited to withstand host changes (13). Similarly, avian generalist *Plasmodium* species may employ analogous strategies, maintaining genetic and epigenetic diversity to adapt rapidly as they shift between avian hosts. However, this remains untested empirically.

Compared to mammalian *Plasmodium* species like *P. falciparum*, avian *Plasmodium* species offer a unique model for examining the strategies of generalist versus specialist parasites. *Plasmodium homocircumflexum* is particularly relevant as it infects multiple host species that are phylogenetically diverse (21). By analyzing gene expression changes in *P. homocircumflexum* across different hosts and over time, we can better understand how parasites navigate the balance between adaptation and specialization. The extent to which transcriptomic changes in generalist parasites stem from immediate host conditions versus retained adaptations from previous hosts remains unclear. It is likely that both factors contribute, as evidenced by *Plasmodium relictum* and *P. homocircumflexum*, which infect a variety of bird species (22). These parasites may rely on multiple transcriptional variation sources to respond effectively to new immune challenges and physiological conditions (23). Identifying the specific sources utilized by *P. homocircumflexum* could enhance our understanding of how generalist parasites optimize their survival across different hosts.

To investigate transcriptional variation, we conducted a crosswise infection experiment where we directly inoculated *P. homocircumflexum* into different host species after the parasite have had time to develop and adjust within either host. The method allowed us to bypass recombination in the vector, ensuring that the parasites initiating the new infection were directly exposed to either the same or a different host environment without time to “reset” their transcriptional profiles. This approach enabled us to assess how the parasite responds to the recipient host and how transcriptional variation unfolds across asexual generations. Different outcomes are expected depending on the underlying mechanism: i) If *P. homocircumflexum* follows a “one key fits all” strategy, gene expression will remain consistent regardless of the host. (ii) If transcriptomic plasticity is the dominant factor, gene expression should reflect the environment of the recipient host, as clonal offspring (i.e. those produced through asexual reproduction) adapt directly to new conditions. (iii) If epigenetic mechanisms drive adaptation, initial gene expression differences influenced by the donor host should gradually adjust to align with the recipient’s conditions. (iv) In cases where selection drives differences, we would expect patterns similar to those seen in the epigenetic scenario, but with increased genetic differentiation (high Fst) among haplotypes, indicating selection on specific variants across generations.

As the infection progresses, we hypothesize that *P. homocircumflexum* may shift gene expression through selection acting on epigenetically diverse asexually reproducing populations or through plastic responses triggered by host immune signals. By examining these scenarios, our study aims to determine whether gene expression in a generalist parasite is primarily shaped by adaptive responses to the recipient host environment or by retained adaptations from previous hosts. This research will deepen our understanding of the adaptability of generalist parasites and the evolutionary pressures that shape host-parasite interactions.

## Results

### 3.1 Parasitemia

Parasitemia from all the individuals changed over the course of the infection (Fig. 1) achieving maximum values around 8 days post infection (dpi) in most of the individuals.

**Figure 1.**
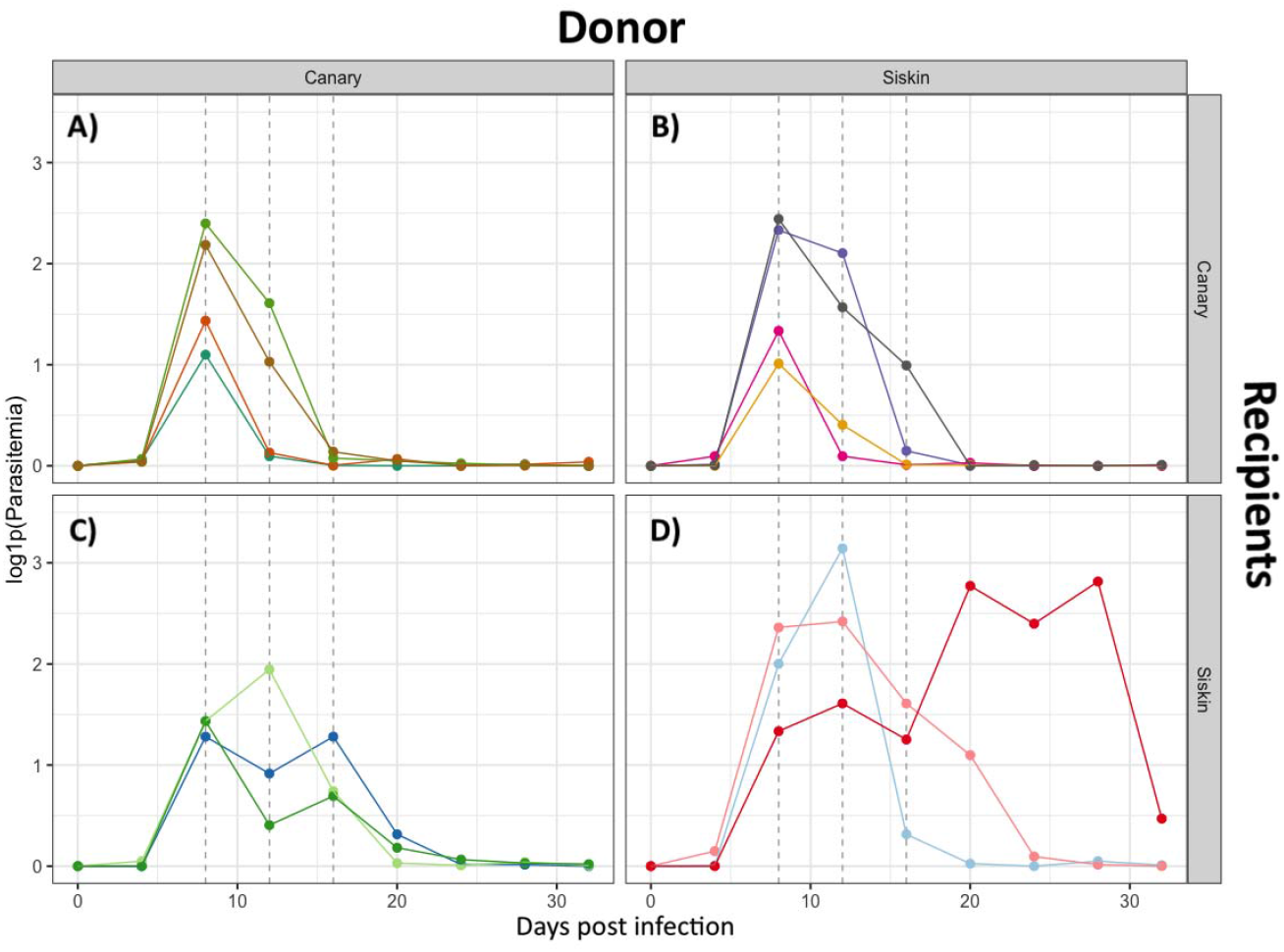
Parasitemia levels for each experimental group. Individuals are colour marked and classified depending on the species donor and species recipient: A) Canaries receiving infected blood from canaries; B) Canaries receiving infected blood from siskins; C) Siskins receiving infected blood from canaries; D) Siskins receiving infected blood from siskin. Dashed lines indicate days of sampling for each group (8, 12 and 16 days post infection).

### 3.2 Principal component analyses

The PCA results obtained from normalized data show distinct clustering patterns for donors and recipients over time (Fig. 2). At 8 dpi, there was a clear separation based on the donor (Fig. 2A), indicating significant differences in their data profiles. However, this separation was not observed when considering the type of recipient (Fig. 2D). At 12 dpi, the distinction between the groups became less pronounced, suggesting a convergence of their data profiles (Fig. 2B-E). At 16 dpi, the PCA plots have shifted to form a tight clustering of samples in the recipient birds of siskins, regardless of the donor (Fig. 2F). In contrast, grouping by donor resulted in more dispersed data points with no clear clustering (Fig. 2).

**Figure 2.**
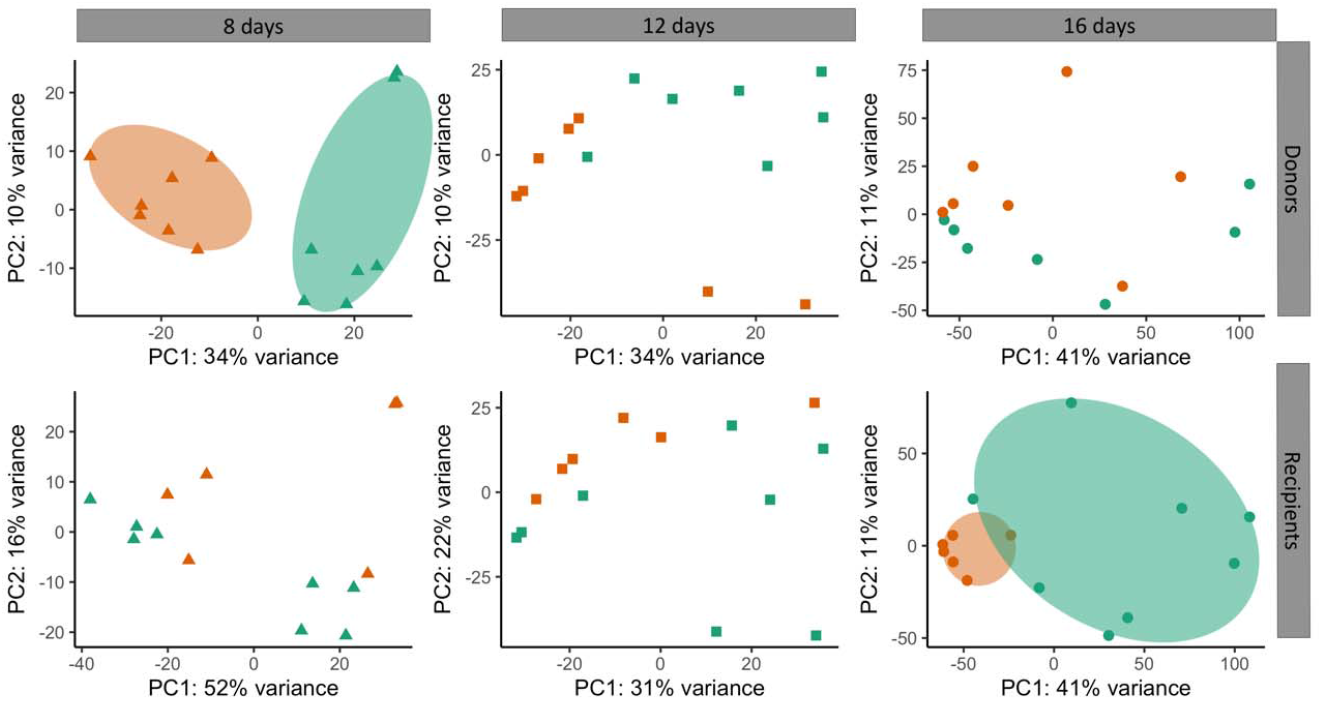
Biplot originating from a Principal Component Analysis (PCA) illustrating the variance in the data for Donors and Recipients at 8, 12, and 16 days post infection (dpi). Shapes represent different time points: triangles indicate 8 dpi, squares 12 dpi, and circles 16 dpi. Colors denote groups, with orange representing siskins and green representing canaries. The first column shows PCA plots of Donors (top row) and Recipients (bottom row) at 8 dpi, the second column at 12 dpi, and the third column at 16 dpi. Each plot displays the percentage of variance explained by the first two principal components (PC1 and PC2). The ellipses represent the 95% confidence intervals for each group.

### 3.3 Differential Expression Analyses

During the course of infection, *Plasmodium homocircumflexum* underwent transcriptional changes as it adapted to the host environment (Fig. 3 and Fig. 4). At 8 and 12 dpi, parasites originating from the same donor showed more homogeneous gene expression patterns, regardless of the species they infected. This observation suggests that early in the infection, parasites relied on pre-established gene expression profiles rather than immediately adapting to the new host environment. This pattern was reflected in the overall expression trends (Fig. 3) and the number of differentially expressed genes (DEGs), which were found to be higher when grouped by donor compared to recipient (Fig. 4). During this phase, parasites may prioritize strategies for immune evasion, red blood cell invasion, and replication while maintaining a transcriptional signature inherited from their original host.

**Figure 3.**
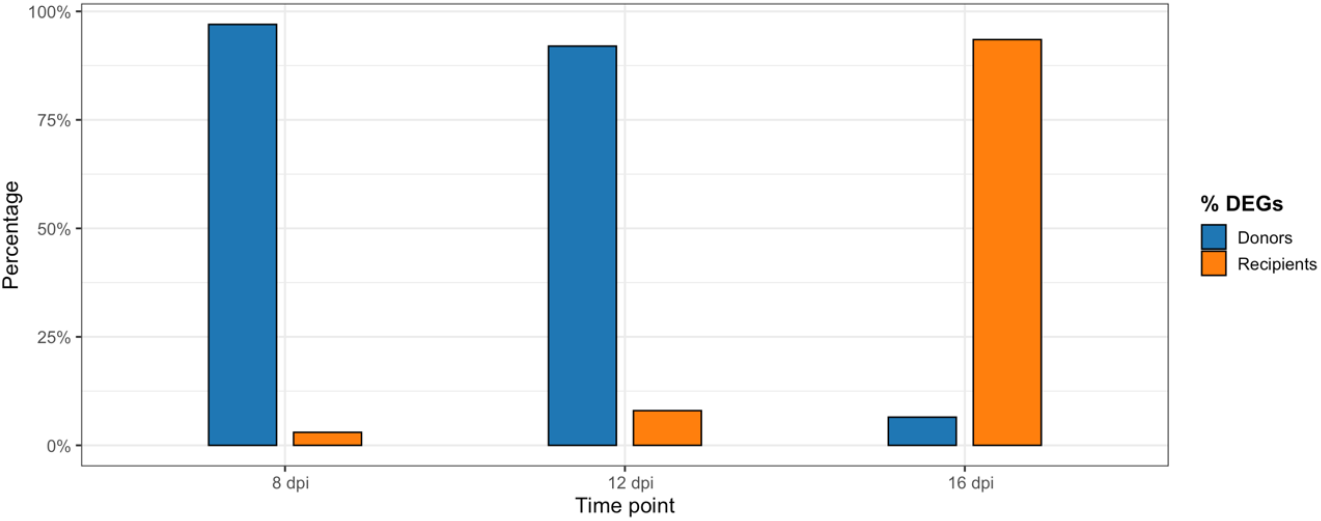
Percentage of differentially expressed genes (DEGs) between donors and recipients at three distinct time points: 8, 12, and 16 days post-infection. For each time point, the percentage of DEGs is represented by two bars, grouped side-by-side for comparison. The blue bars correspond to donors, while the orange bars represent recipients.

**Figure 4.**
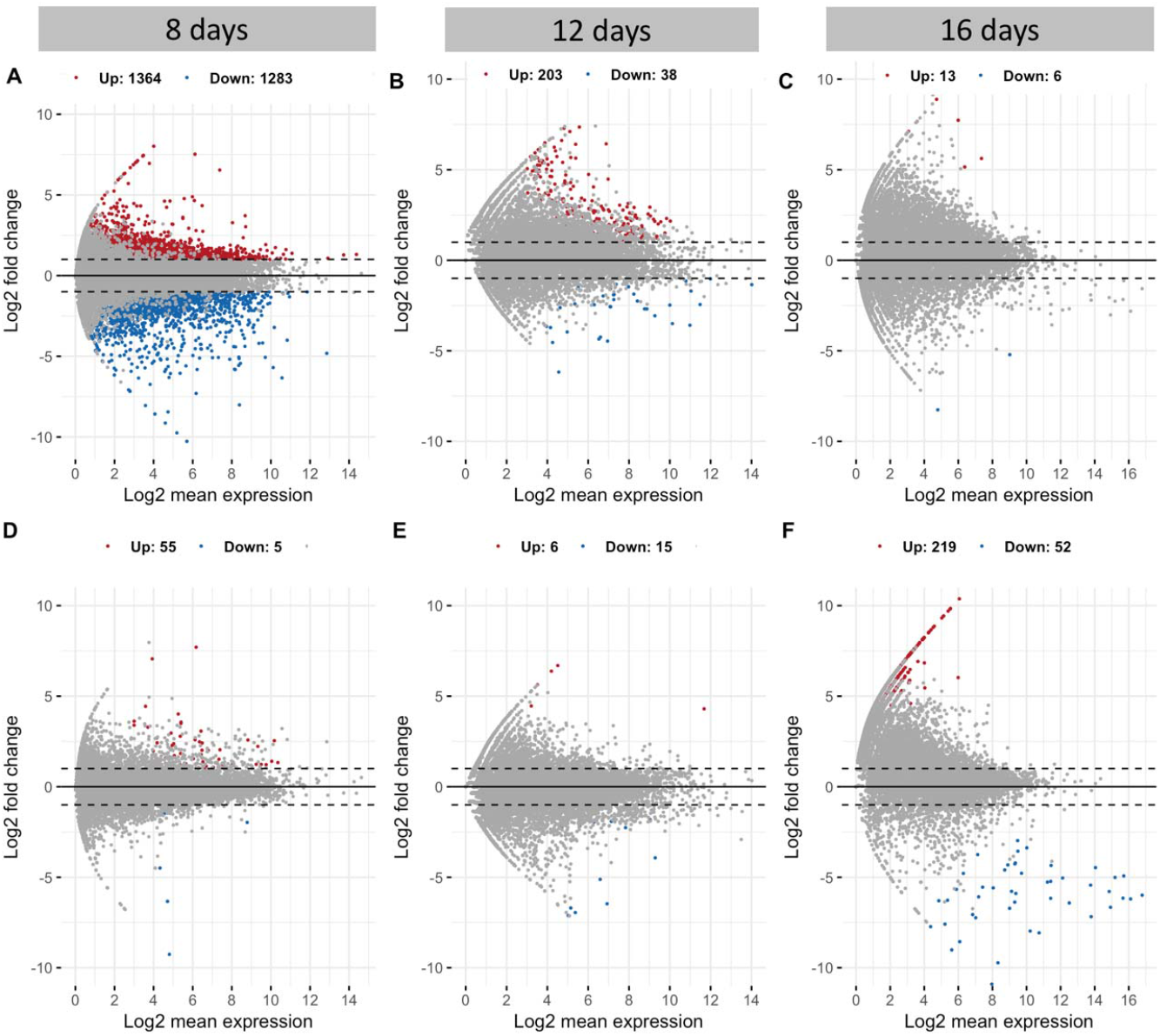
MA plots showing differential gene expression analysis at 8, 12 and 16 days post infection. The three plots on the top (A,B,C) show the results when analyzing the data by taking into account the donor species. The three plots on the bottom (D,E,F) show the results when analyzing the data by taking into account the recipient species. The x-axis represents the log2 fold change, and the y-axis shows the −log10 adjusted p-value. Red points indicate significantly upregulated genes, blue points indicate significantly downregulated genes, and gray points are non-significant. Highlighted gene names are the top 20 differentially expressed genes (DEGs) (fdr < 0.05, |log2FoldChange| > 2). Also, the number of DEGs are indicated in numbers for each plot.

However, by 16 dpi, the influence of the recipient species emerged as the primary factor shaping parasite gene expression (Fig. 3). At this stage, transcriptomic profiles were more distinct between host species, suggesting an active response to new physiological and immunological pressures. DEG patterns also shifted, with a higher percentage of DEGs associated with the recipient rather than the donor (Fig. 4). The most significant changes occurred between 8 dpi and 16 dpi. At 8 dpi, 2,647 DEGs were detected when grouping by donors (Fig. 4A), while only 60 DEGs were observed when considering recipients (Fig. 4D). By 16 dpi, this trend reversed, with 271 DEGs when grouping by recipients (Fig. 4F) and only 19 DEGs by donors (Fig. 4C). These findings suggest that as infection progressed, parasites shifted from a generalist gene expression strategy to a more specialized adaptation tailored to the recipient host, likely involving metabolic adjustments and mechanisms for immune evasion. This highlights the plasticity of *Plasmodium homocircumflexum* in modulating its transcriptome to optimize survival across different host environments.

### 3.4 Variant calling

A distribution of the samples according to their SNPs was obtained for day 8 post infection (Fig 5). As indicated in the materials and methods section, similar analysis could not be performed for 12 and 16 dpi. The results indicate that samples do not cluster clearly when classified according to either recipient (Fig 5A) or donor (Fig 5B) species. The distribution of the samples showed no clear pattern when considering either the recipient (Fig 5A) or donor (Fig 5B) species. Additionally, two FST analyses were conducted. The first analysis yielded an FST value of 0.09 (95% CI 0.02 - 0.15) when considering donor species, while the second analyses resulted in an FST of 0.03 (95% CI 0.01 - 0.14) when considering recipient species. These FST values further support a lack of clear genetic differentiation among the samples based on the species classification.

**Figure 5.**
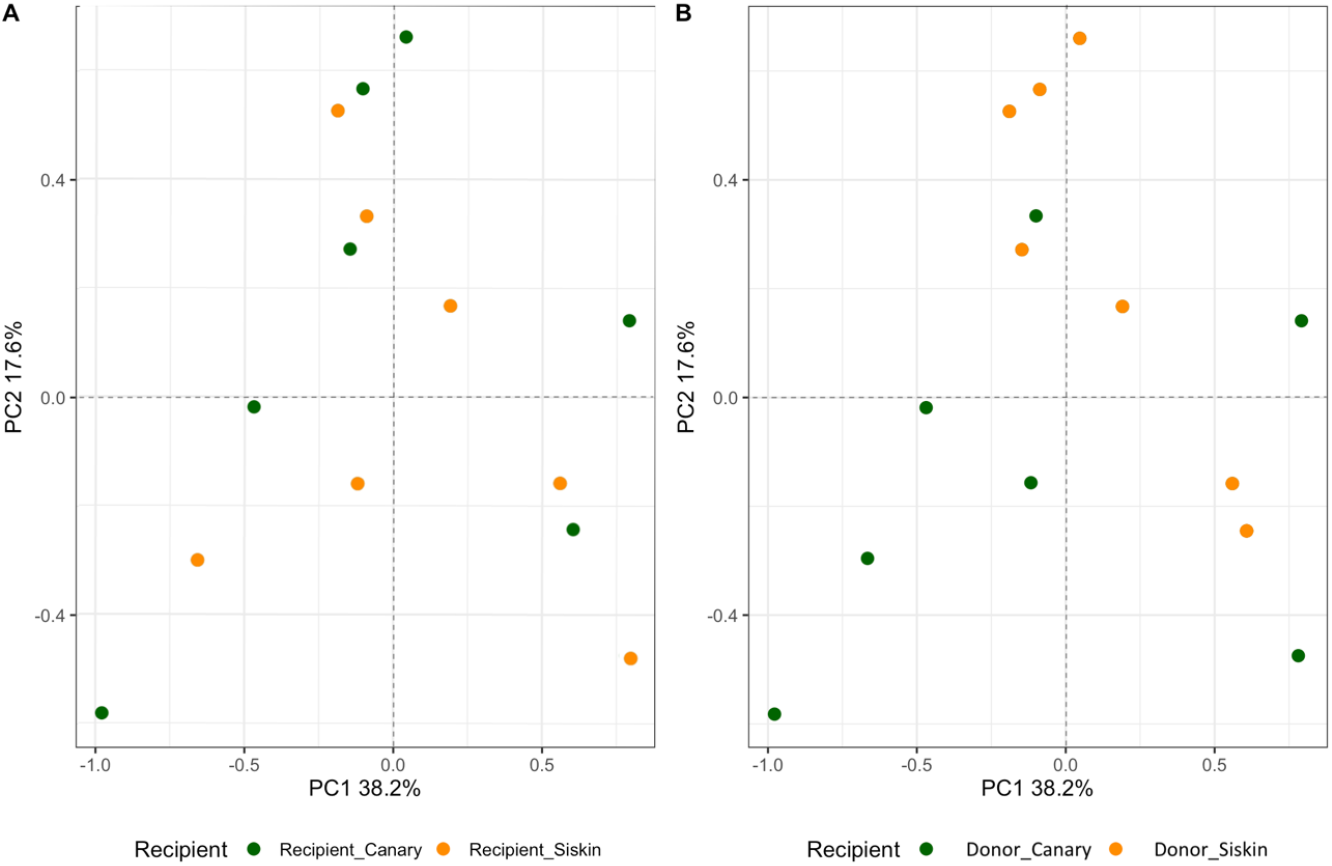
PCA of haploid samples from 8dpi classified based on two different criteria. Panel (A) represents the distribution of samples when classified according to the recipient species. Panel (B) shows the distribution of the same samples but classified based on the blood donor.

## Discussion

Our results provide insights into *P. homocircumflexum* infection in two avian host species, showing how generalist parasites adapt to phylogenetically distinct hosts. Parasitemia peaked at 8 dpi, with RNA expression shifting from donor-dependent at 8 dpi to recipient-dependent by 16 dpi. Regarding the scenarios: (i) The “one key fits all” strategy could be ruled out, as gene expression shifted according to the host environment. (ii) Transcriptomic plasticity could be partially supported, as expression in later stages reflected the recipient host environment, suggesting adaptation. (iii) An epigenetic mechanism could be supported, with initial expression differences influenced by the donor host regardless of recipient host, followed by a gradual adjustment to the recipient’s conditions. (iv) Selection-driven differences is not supported, as no elevated genetic differentiation was observed in the SNP analyses. These findings underscore the plasticity and adaptability of generalist parasite plasticity in diverse host environments. Below we discuss these results in detail.

Although the infection dynamics of some parasites are well understood, classifying a parasite as a generalist or specialist depends on the density of its reservoir host (8). In this context, the biodiversity and abundance of species within an ecosystem may influence whether a parasite lineage is more or less generalist in its host range (1, 25, 26). Indeed, previous studies have determined that the same parasite species can adapt to a more specialist role in environments with a limited number of potential host species, compared to scenarios where many potential hosts are available (27, 28). The molecular mechanisms and adaptations that a generalist parasite population can undergo during infection and reproduction in different host species have been insufficiently analyzed. Our results provide new insights into this issue. Despite exhibiting similar levels of parasitemia, we revealed that the source of infection significantly influences the RNA expression patterns of the parasite. In essence, the donor species seems to dictate parasite expression in the days following infection. However, after 16 dpi, expression patterns appear to be primarily determined by the infected species (i.e., the recipient). According to the existing scientific literature, factors contributing to these changes in RNA expression patterns may include genetic variations such as SNPs, epigenetic modifications, or direct plasticity (18).

Single nucleotide polymorphisms in human malaria have been pivotal in identifying genetic factors that influence susceptibility to infection and drug resistance (29, 30). Over the last decade, SNP studies have been extensively used to uncover host-pathogen interactions and guide the development of targeted therapies and vaccines (e.g. 31). In this study, we have used established SNP analysis workflows to determine whether the differences found during peak infection (8 dpi) could be due to natural selection within the parasite population. Our findings indicate that genetic selection during peak infection is 9% based on donor species and 3% based on recipient species. Although these percentages suggest a small degree of selection at this stage of the infection cycle, we cannot definitively conclude that this process accounts for the observed differences.

Epigenetic mechanisms, such as histone modifications and DNA methylation, have been recognized as significant drivers of transcriptional variation in parasites. These mechanisms enable heritable and reversible changes in chromatin state, which are crucial for the adaptive capacity of these organisms (32, 33). This flexibility is particularly important for generalist parasites, as it facilitates their adaptation to changing environments during infection (12, 34). In mammalian malaria systems, including *P. falciparum*, epigenetic factors have been shown to drive early gene expression changes (19). It is plausible that *P. homocircumflexum* employs similar mechanisms inherited from the donor population to regulate gene expression at 8 dpi. These inherited epigenetic responses may provide a strategic advantage during the early stages of infection, allowing for rapid adaptation to the new host environment. As the infection progresses, the host immune system could exert selective pressure (35), resulting in recipient-dependent patterns of gene expression. This shift underscores the parasite’s dynamic adaptability in response to host-derived signals and immune challenges, highlighting the complexity of generalist parasite-host interactions. To test this hypothesis, a detailed comparison of RNA expression profiles between the source population (donors) and the parasite population at 8 dpi would be necessary. Such analyses could clarify whether transcriptional patterns reflect inherited epigenetic regulation or other adaptive processes.

Direct transcriptional variation refers to transient, non-heritable changes in gene expression that are triggered by environmental stressors (18). In the context of host-parasite interactions, these mechanisms can be influenced by the host immune response, which evolves dynamically over the course of an infection. Around 12 dpi, the host immune system transitions from innate to adaptive immunity (36, 37), marking a critical turning point in the interaction between the host and the parasite. In our study, we observed a distinct shift in RNA expression patterns between 8 and 16 dpi. Early in the infection (8 dpi), parasite gene expression appeared to be dependent on the donor population. However, by 16 dpi, it was predominantly shaped by the recipient species. This temporal change aligns with the timing of immune system modulation, as the influence of the donor diminishes and the adaptive immune signals from the recipient take precedence. Our findings support the idea that parasite transcriptional responses are not only reactive but also adaptive to the host immune environment. This highlights the role of immune system dynamics in shaping gene expression. These results provide valuable insights into how immune responses drive transcriptional variation, furthering our understanding of the complex interplay between host and parasite.

A critical next step is to investigate the host immune response over time, with a particular focus on distinct immune pathways or gene groups that are activated at different stages of infection. RNA-seq analyses could be instrumental in uncovering pathways associated with innate immunity during the early stages and adaptive immunity in the later stages of infection, thereby providing valuable insights into host-parasite dynamics. Furthermore, exploring cytokine responses, T-cell activation pathways, and other immune-related processes would help clarify the role of host immunity in shaping parasite gene expression. Such analyses could elucidate whether specific pathways are consistently targeted by *P. homocircumflexum* across hosts, which would improve our understanding of host-specific versus generalist responses.

Our experimental design, which involved blood inoculations, ensured that parasite populations began from the same starting point. This controlled setup allowed us to detect significant differences in RNA profiles influenced by donor species at 8 dpi and by recipient species at 16 dpi. The donor effect observed during early stages highlights the importance of parasite synchronization with the donor host. These results suggest that gene expression patterns may differ in natural infections transmitted by vectors due to the additional epigenetic regulation imposed by vector stages. Follow-up studies that include vector-host-parasite interactions would provide valuable insights into RNA expression dynamics and the influence of epigenetic regulation during transmission. Moreover, comparisons with human malaria experiments, where vector-stage regulation has been well documented (38), could further refine our understanding of these processes in avian systems.

In conclusion, our study highlights the dynamic transcriptional plasticity of *P. homocircumflexum* during infection in two avian host species, emphasizing how generalist parasites respond to distinct host environments. We observed that parasitemia peaked at 8 dpi, with RNA expression patterns initially influenced by the donor species before shifting to profiles driven by the recipient by 16 dpi. This transition may coincide with the progression of the host immune response from innate to adaptive immunity, suggesting that host-derived signals could influence parasite transcriptional responses. However, since we did not directly assess host immune markers, further studies are needed to confirm this relationship. Our SNP analyses revealed limited genetic selection, indicating that the observed transcriptional shifts are unlikely driven by genetic changes but may instead result from epigenetic mechanisms such as histone modifications or DNA methylation. These mechanisms may provide the parasite with transcriptional flexibility necessary to rapidly adjust to host-specific environments. Nonetheless, whether these changes are adaptive in the context of transmission remains unclear. In particular, the relationship between within-host transcriptional shifts and transmission success warrants further investigation. If the genes undergoing regulation are involved in gametocyte production, this could indicate an adaptive response facilitating transmission. Future research incorporating gametocyte-stage expression data and vector-mediated infections will be crucial to fully understand the interplay between host immunity, epigenetic regulation, and parasite adaptation.

## Materials and Methods

### 2.1 Experimental animals and parasite acquisition

The research was conducted in 2019 at the Nature Research Centre in Vilnius, Lithuania, using juvenile domestic canaries (*Serinus canaria domestica*) and Eurasian siskins (*Spinus spinus*) from commercial suppliers. All procedures complied with European and Lithuanian regulations on animal research, with ethical approval from the State Food and Veterinary Service of Lithuania (Approval No. 2018/05/03-G2-84).

Birds were housed individually in cages, allowing social interaction, in a vector-free room with a stable 21 ± 1 °C temperature and a natural light/dark cycle. Food and water were provided *ad libitum*. Before experiments, PCR and microscopy confirmed all birds were malaria-free. The parasite used was *Plasmodium* (Giovannolaia) *homocircumflexum*, lineage COLL4 (GenBank KC884250), originally isolated in 2010 from a wild Red-backed shrike (*Lanius collurio*) captured at the Biological Station of the Zoological Institute of the Russian Academy of Sciences on the Curonian Spit, Baltic Sea. The strain was cryopreserved following (39) and later used for experiments.

## 2.2 Experimental procedures

Eight canaries and six siskins were randomly assigned to four experimental groups (Fig. 6). The *P. homocircumflexum* strain COLL4, obtained from a canary, was used to infect four canaries and three siskins. The same strain, sourced from a siskin, was used to infect four additional canaries and three siskins. The experiments used the parasite’s seventh passage.

**Figure 6.**
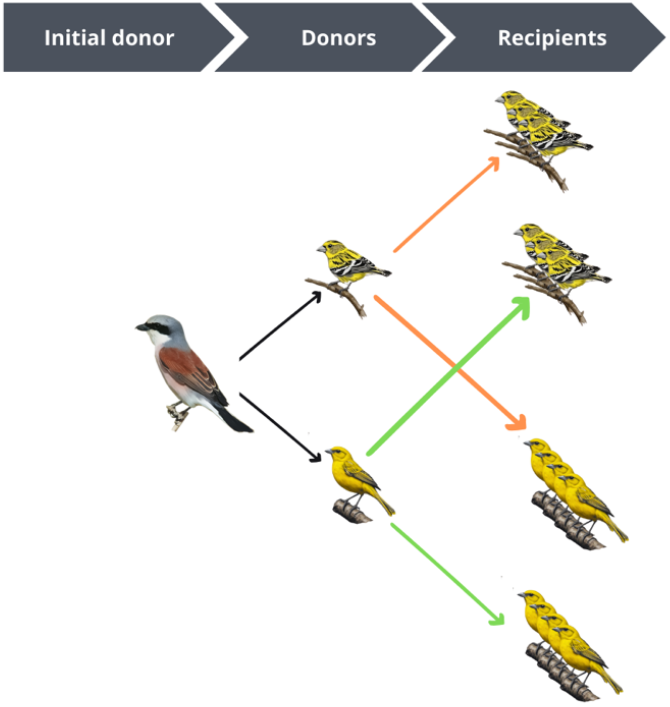
Experimental design for this study involved using blood from an initial donor for consecutive experimental infections. One canary and one siskin served as blood donors, while a total of 6 siskins and 8 canaries were used as recipients. Colored arrows indicate the origin of the infected blood: orange arrows represent blood from siskin donor, while green arrows represent blood from canary donor. (Drawings obtained with the artificial intelligence of artbreeder and the final figure designed with Canva)

To initiate infections, cryopreserved blood was introduced into a non-infected canary and siskin, which then served as donors: canary-derived parasites for canaries and siskins, and siskin-derived parasites for canaries and siskins. Each bird received 100 µL of a freshly prepared suspension containing infected blood, 3.7% sodium citrate, and 0.9% saline (4:1:5 ratio), with an erythrocytic meront intensity of 0.4% and an estimated dose of 6×10^5^ meronts. The mixture was injected into the pectoral muscle following Iezhova et al. (40). Birds were monitored for 32 dpi, with blood samples collected every four days for microscopy and PCR. Additional samples for RNA sequencing were taken at 8, 16, and 24 dpi.

### 2.3 Sampling, Microscopy and Molecular Analysis

Blood samples were obtained by puncturing the brachial vein (41). A small drop of blood from each bird was used to prepare two blood smears, while approximately 30 µl was stored in SET-buffer (0.05 M Tris, 0.15 M NaCl, 0.5 EDTA, pH 8.0) for molecular analysis Hellgren et al. (42). Samples for RNA analysis were preserved in TRIzol LS Reagent (Invitrogen, Carlsbad, CA). Blood samples were kept at −20°C until processed. The blood smears were air-dried, fixed with absolute methanol, and stained with Giemsa (43). The slides were examined under an Olympus BX51 light microscope equipped with an Olympus DP12 (Olympus, Shinjuku City, Japan). Approximately 100 fields were reviewed at a magnification of 1000×. Parasitemia was quantified by counting the number of parasites per 1,000 erythrocytes, or per 10,000 erythrocytes in cases of low infection intensity, following the protocol recommended by Godfrey et al. (44). Detailed procedures for preparing, staining, and examining blood smears were outlined by Valkiūnas et al. (43).

DNA extraction was performed using the ammonium-acetate protocol (45). A nested-PCR protocol was employed for genetic analysis (46). Amplification products (1.5 µl) were resolved on a 2% agarose gel. Sequencing followed the protocol by Bensch et al. (47), targeting the 5’ region with the primer HaemF. Dye terminator cycle sequencing (BigDye) was conducted on an ABI PRISMTM 3100 capillary sequencer (Applied Biosystems, USA). Sequence editing and alignment were carried out using BioEdit (48).

### 2.4 RNA extraction and sequencing

Blood samples for RNA sequencing were collected at three different time points (8 dpi, 16 dpi and 12 dpi). The samples were stored in TRIzol. Total RNA was extracted from 20 μL whole blood using 1000 μL TRIzol LS Reagent (Invitrogen, Carlsbad, CA) and homogenised by vortexing from all seven individuals. Samples were then incubated at room temperature for 5 min before the addition of 200 μl chloroform (Merck KGaA, Darmstadt, Germany). After a further incubation at room temperature for 3 min, the samples were centrifuged at 11,000 rpm for 17 min at 4°C. The supernatant was then transferred to new tubes and processed using a RNeasy Mini Kit (Qiagen, GmbH, Hilden, Germany). Following the manufacturer’s protocol, 1 volume of 70% ethanol was added to the lysate. The total extracted RNA was shipped on dry ice to Novogene Bioinformatics Technology, Hong Kong, for RNA quality control, DNAse treatment, rRNA reduction and amplification using the SMARTer Ultra Low Kit (Clontech Laboratories, Inc.). Novogene performed library preparation, cDNA synthesis and paired-end RNA sequencing using the Illumina HiSeq 2000. We quality checked all demultiplexed RNA-seq reads using FastQC (v.0.10.1).

### 2.5 De novo transcriptome assembly

As the genome of *P. homocircumflexum* is not available, alternative tools were employed in order to isolate information belonging exclusively to the *P. homocircumflexum* lineage pCOLL4. Initially, the canary genome (NCBI RefSeq assembly serCan2020) and a portion of the siskin genome (NCBI RefSeq assembly ASM3478079v1) were employed to eliminate reads belonging to the bird. Subsequently, the P*lasmodium relictum* genome (NCBI RefSeq assembly GCA_900005765.1; 49) was utilized to keep reads belonging to the malaria parasite. Finally, the *P. homocircumflexum* transcriptome previously published (50) was employed to obtain a greater number of reads and ensure that the reads used exclusively belonged to the malaria parasite. Analyses with and without the reference genome were performed using the bioinformatics software STAR (51) and Bowtie (52). Standard adjustments were implemented to determine alignment sensitivity. Finally, the de novo assembly of the parasite was conducted using the transcriptome assembler Trinity (v. 2.15.1) (53), resulting in 14 649 contigs.

GC content varies across different eukaryotic organisms, with where *Plasmodium* species evolving toward a high AT richness compared to its hosts (54, 55, 56, 57), which provides a valuable tool for transcript separation in host-parasite studies. Consequently, the subsequent transcript was subjected to filtration based on GC content in order to exclude any host contigs and to include sequences with a mean GC content below 23% (n = 12 058).

### 2.6 Differential gene expression level

The expression levels of the parasite genes were quantified using Salmon (v. 1.3.2) (58). Salmon is a software program that produces expected read counts for every contig and identifies genes that are expressed between species, along with the name of each contig. The read counts for every contig were stored in a file that was statistically analyzed inside the R statistical environment (v. 4.4.0) (59). Read counts were normalized using regularized log transformation in order to account for potential variation in sequencing depth and the large differences in the number of parasites present in the blood (parasitaemia levels). Regularized log transformation of counts was performed in order to represent the data without any prior knowledge of the sampling design in the principal component analysis (PCA) and sample distance calculations. This method of presenting counts without bias is preferable to variance stabilizing of counts when the size factors vary greatly between the samples, as is the case in our data. The package ggplot2 (60) was employed for the generation of all graphs.

### 2.7 Statistical analyses

For the differential expression analyses we used the R package DeSeq2 (61), which is designed for the specific task of performing this type of analysis. The DeSeq2 package (version 1.16.1) (61) was utilized to correct the variance-mean dependence in count data derived from high-throughput sequencing assays and to test for differential expression based on a model that employs the negative binomial distribution. When significant differences in expression were identified, and to circumvent potential issues associated with sequencing depth, gene length, or RNA composition, count data were initially normalised using the DeSeq2 method.

The statistical comparisons are presented in Fig 6. Firstly, the samples were classified according to the donor. This distinction was made regardless of the species of bird receiving the blood (Fig 6). Subsequently, the samples were classified based on the recipient of the blood (Fig 6). In this instance, only the species of bird receiving the blood was considered. This approach enabled the number of differentially expressed genes (DEGs) to be determined in relation to either the origin of the blood (from birds of mixed species but with the same donor species) or the recipient (from birds of the same species but with infection originating from different species of donor).

## Acknowledgments

Funding was provided by the Junta de Extremadura (PO17024, Post-Doc grant) to LGL, the Spanish Ministry of Science and Innovation (PID2022-140397NB-I00) to AM, and LA4 (R+D+I program in the Biodiversity Area financed with the funds of the FEDER Extremadura 2021–2027 Operational Program of the Recovery, Transformation and Resilience Plan) to LGL and AM. VP obtained funding from the Research Council of Lithuania (Project No. 09.3.3-LMT-K-712-01-0016). OH obtained funding from the Swedish research council (VR 2016-03419 and 2021-03663). All the authors would like to thanks to Ananias Escalante for taking the time to read the manuscript and for his inspiring ideas.

